# OrthoSNAP: a tree splitting and pruning algorithm for retrieving single-copy orthologs from gene family trees

**DOI:** 10.1101/2021.10.30.466607

**Authors:** Jacob L. Steenwyk, Dayna C. Goltz, Thomas J. Buida, Yuanning Li, Xing-Xing Shen, Antonis Rokas

## Abstract

Molecular evolution studies, such as phylogenomic studies and genome-wide surveys of selection, often rely on gene families of single-copy orthologs (SC-OGs). Large gene families with multiple homologs in one or more species—a phenomenon observed among several important families of genes such as transporters and transcription factors—are often ignored because identifying and retrieving SC-OGs nested within them is challenging. To address this issue and increase the number of markers used in molecular evolution studies, we developed OrthoSNAP, a software that uses a phylogenetic framework to simultaneously split gene families into SC-OGs and prune species-specific inparalogs. We term SC-OGs identified by OrthoSNAP as SNAP-OGs because they are identified using a *<u>s</u>*plitti*<u>n</u>*g *<u>a</u>*nd *<u>p</u>*runing procedure analogous to snapping branches on a tree. From 415,129 orthologous groups of genes inferred across seven eukaryotic phylogenomic datasets, we identified 9,821 SC-OGs; using OrthoSNAP on the remaining 405,308 orthologous groups of genes, we identified an additional 10,704 SNAP-OGs. Comparison of SNAP-OGs and SC-OGs revealed that their phylogenetic information content was similar, even in complex datasets that contain a whole genome duplication, complex patterns of duplication and loss, transcriptome data where each gene typically has multiple transcripts, and contentious branches in the tree of life. OrthoSNAP is useful for increasing the number of markers used in molecular evolution data matrices, a critical step for robustly inferring and exploring the tree of life.

## Introduction

Molecular evolution studies, such as species tree inference, genome-wide surveys of selection, evolutionary rate estimation, measures of gene-gene coevolution, and others typically rely on single-copy orthologs (SC-OGs), a group of homologous genes that originated via speciation and are present in single-copy among species of interest [1–6]. In contrast, paralogs—homologous genes that originated via duplication and are often members of large gene families—are typically absent from these studies (Fig 1). Gene families of orthologs and paralogs often encode functionally significant proteins such as transcription factors, transporters, and olfactory receptors [7–10]. The exclusion of SC-OGs from gene families has not only hindered our understanding of their evolution and phylogenetic informativeness but is also artificially reducing the number of gene markers available for molecular evolution studies. Furthermore, as the number of species and / or their evolutionary divergence increases in a dataset, the number of SC-OGs decreases [11,12]; case in point, no SC-OGs were identified in a dataset of 42 plants [11]. As the number of available genomes across the tree of life continues to increase, our ability to identify SC-OGs present in many taxa will become more challenging.

**Figure 1.**
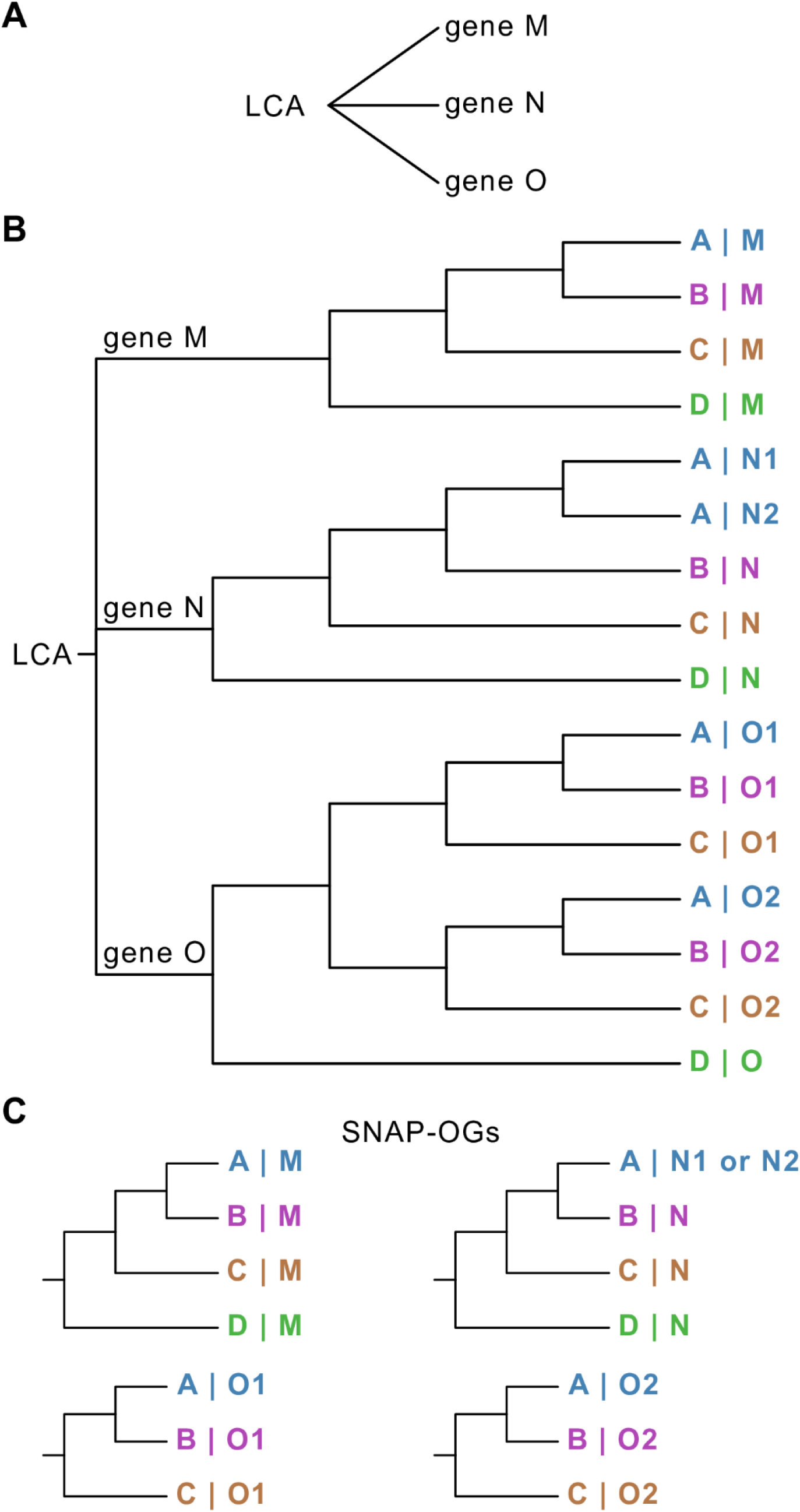
Cartoon depiction of three classes of paralogs: outparalogs, inparalogs, and coorthologs. (A) Paralogs refer to related genes that have originated via gene duplication, such as genes M, N, and O. (B) Outparalogs and inparalogs refer to paralogs that are related to one another via a duplication event that took place prior to or after a speciation event, respectively. With respect to the speciation event that led to the split of taxa A, B, and C from D, genes M, N, and O are outparalogs because they arose prior to the speciation event; genes O1 and O2 in taxa A, B, and C are inparalogs because they arose after the speciation event. Species-specific inparalogs are paralogous genes observed only in one species, strain, or organism in a dataset, such as gene N1 and N2 in species A. Species-specific inparalogs N1 and N2 in species A are also coorthologs of gene N in taxa B, C, and D; the same is true for inparalogs O1 and O2 from species A, which are coorthologs of gene O from species D. (C) Cartoon depiction of SNAP-OGs identified by OrthoSNAP.

In light of these issues, several methods have been developed to account for paralogs in specific types of molecular evolution studies—for example, in species tree reconstruction [13]. Methods such as SpeciesRax, STAG, ASTRAL-PRO, and DISCO can be used to infer a species tree from a set of SC-OGs and gene families composed of orthologs and paralogs [11,14–16]. Other methods such as PHYLDOG [17] and guenomu [18] jointly infer the species and gene trees, but require abundant computational resources, which has hindered their use for large datasets. Other software, such as PhyloTreePruner, can conduct species-specific inparalog trimming [19]. Agalma, as part of a larger automated phylogenomic workflow, can prune gene trees into maximally inclusive subtrees wherein each species, strain, or organism is represented by one sequence [20]. Similarly, OMA identifies subgroups of single-copy orthologs using graph-based clustering of sequence similarity scores [21]. Although these methods have expanded the numbers of gene markers used in species tree reconstruction, they were not designed to facilitate the retrieval of as broad a set of SC-OGs as possible for downstream molecular evolution studies such as surveys of selection. Furthermore, the phylogenetic information content of these gene families remains unknown, calling into question their usefulness.

To address this need and measure the information content of subgroups of single-copy orthologous genes, we developed OrthoSNAP, a novel algorithm that identifies SC-OGs nested within larger gene families via tree decomposition and species-specific inparalog pruning. We term SC-OGs identified by OrthoSNAP as SNAP-OGs because they were retrieved using a *<u>s</u>*plitti*<u>n</u>*g *<u>a</u>*nd *<u>p</u>*runing procedure. The efficacy of OrthoSNAP and the information content of SNAP-OGs was examined across seven eukaryotic datasets, which include species with complex evolutionary histories (e.g., whole genome duplication) or complex gene sequence data (e.g., transcriptomes, which typically have multiple transcripts per protein-coding gene). These analyses revealed OrthoSNAP can substantially increase the number of orthologs for downstream analyses such as phylogenomics and surveys of selection. Furthermore, we found that the information content of SNAP-OGs was statistically indistinguishable from that of SC-OGs suggesting the inclusion of SNAP-OGs in downstream analyses is likely to be as informative. These analyses indicate that SNAP-OGs identified by OrthoSNAP hold promise for increasing the number of markers used in molecular evolution studies, which can, in turn, be used for constructing and interpreting the tree of life.

## Results

OrthoSNAP is a novel tree traversal algorithm that conducts tree splitting and species-specific inparalog pruning to identify SC-OGs nested within larger gene families (Fig. 1C). OrthoSNAP takes as input a gene family phylogeny and associated FASTA file and can output individual FASTA files populated with sequences from SNAP-OGs as well as the associated Newick tree files (Fig. 2). During tree traversal, tree uncertainty can be accounted for by OrthoSNAP by collapsing poorly supported branches. In a set of seven eukaryotic datasets that contained 9,821 SC-OGs, we used OrthoSNAP to identify an additional 10,704 SNAP-OGs. Using a combination of multivariate statistics and phylogenetic measures, we demonstrate that SNAP-OGs and SC-OGs have similar phylogenetic information content in all seven datasets. This observation was consistent across datasets where the identification of large numbers of SC-OGs is challenging: flowering plants that have complex patterns of gene duplication and loss (15 SC-OGs and 653 SNAP-OGs), a lineage of budding yeasts wherein half of the species have undergone an ancient whole-genome duplication event (2,782 SC-OGs and 1,334 SNAP-OGs), and a dataset of transcriptomes where many genes are represented by multiple transcripts (390 SC-OGs and 2,087 SNAP-OGs). Lastly, similar patterns of support were observed among the 252 SC-OGs and the 1,428 SNAP-OGs in a contentious branch in the tree of life. Taken together, these results suggest that OrthoSNAP is helpful for expanding the set of gene markers available for molecular evolutionary studies, even in datasets where inference of orthology has historically been difficult due to complex evolutionary history or complex data characteristics.

**Figure 2.**
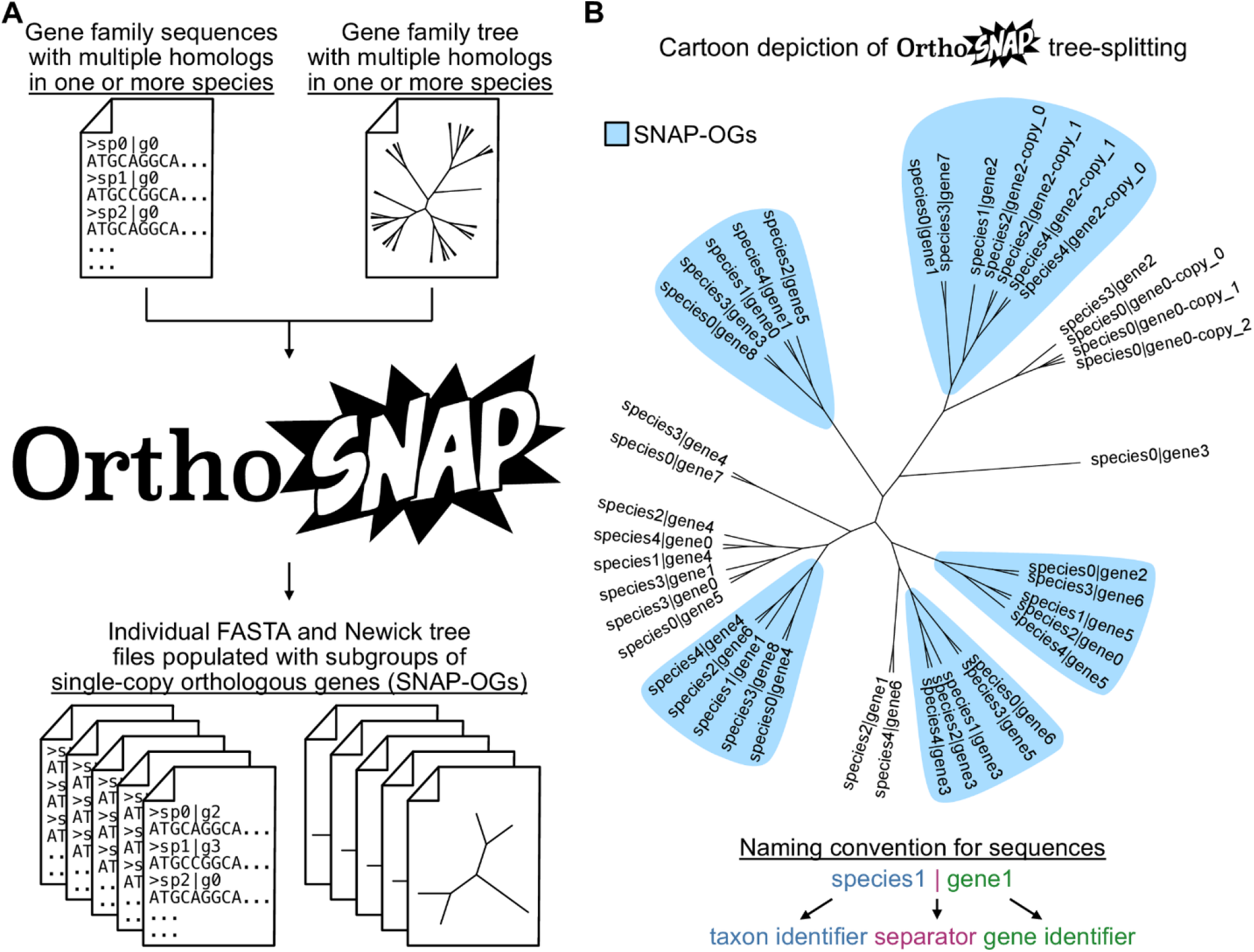
Cartoon depiction of OrthoSNAP workflow. (A) OrthoSNAP takes as input two files: a FASTA file of a gene family with multiple homologs observed in one or more species and the associated gene family tree. The outputted file(s) will be individual FASTA files of SNAP-OGs. Depending on user arguments, individual Newick tree files can also be outputted. (B) A cartoon phylogenetic tree that depicts the evolutionary history of a gene family and five SNAP-OGs therein. While identifying SNAP-OGs, OrthoSNAP also identifies and prunes species-specific inparalogs (e.g., species2|gene2-copy_0 and species2|gene2-copy_1), retaining only the inparalog with the longest sequence, a practice common in transcriptomics. Note, OrthoSNAP requires that sequence naming schemes must be the same in both sequences and follow the convention in which a species, strain, or organism identifier and gene identifier are separated by pipe (or vertical bar; “|”) character.

### SC-OGs and SNAP-OGs have similar information content

To compare SC-OGs and SNAP-OGs, we first independently inferred orthologous groups of genes in three eukaryotic datasets of 24 budding yeasts (none of which have undergone whole-genome duplication), 36 filamentous fungi (*Aspergillus* and *Penicillium* species), and 26 mammals including humans, dogs, pigs, elephants, sloths, and others (Table S1). There was variation in the number of SC-OGs and SNAP-OGs in each lineage (S1 Fig; Table S2). Interestingly, the ratio of SNAP-OGs : SC-OGs among budding yeasts, filamentous fungi, and mammals was 0.83 (1,392 : 1,668), 0.46 (2,035 : 4,393), and 5.53 (1,775 : 321), respectively, indicating SNAP-OGs can substantially increase the number of gene markers in certain lineages. The number of SNAP-OGs identified in a gene family with multiple homologs in one or more species also varied (S2 Fig).

Similar orthogroup occupancy and best fitting models of substitutions were observed among SC-OGs and SNAP-OGs (S3 Fig; Table S3), raising the question of whether SC-OGs and SNAP-OGs have similar information content. To answer this, the information content among multiple sequence alignments and phylogenetic trees from SC-OGs and SNAP-OGs (S4 Fig; Table S4) was compared across nine properties—Robinson-Foulds distance [22], relative composition variability [23], and average bootstrap support, for example—using multivariate analysis and statistics as well as information theory-based phylogenetic measures. Principal component analysis enabled qualitative comparisons between SC-OGs and SNAP-OGs in reduced dimensional space and revealed a high degree of similarity (Fig 3, S5 Fig). Multivariate statistics—namely, multi-factor analysis of variance—facilitated a quantitative comparison of SC-OGs and SNAP-OGs and revealed no difference between SC-OGs and SNAP-OGs (p = 0.63, F = 0.23, df = 1; Table S5) and no interaction between the nine properties and SC-OGs and SNAP-OGs (p = 0.16, F = 1.46, df = 8). Similarly, multi-factor analysis of variance using an additive model, which assumes each factor is independent and there are no interactions (as observed here), also revealed no differences between SC-OGs and SNAP-OGs (p = 0.65, F = 0.21, df = 1). Next, we calculated tree certainty, an information theory-based measure of tree congruence from a set of gene trees, and found similar levels of congruence among phylogenetic trees inferred from SC-OGs and SNAP-OGs (Table S6). Taken together, these analyses demonstrate that SC-OGs and SNAP-OGs have similar phylogenetic information content.

**Figure 3.**
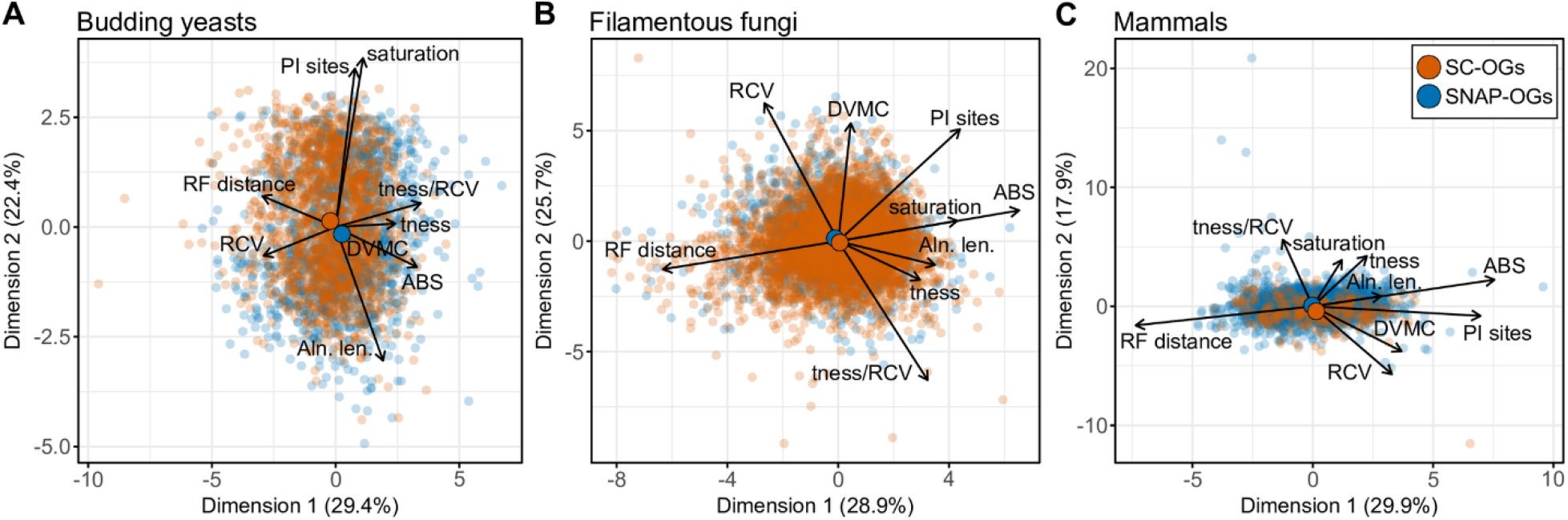
SC-OGs and SNAP-OGs have similar phylogenetic information content. To evaluate similarities and differences between SC-OGs (orange dots) and SNAP-OGs (blue dots), we examined each gene’s phylogenetic information content by measuring nine properties of multiple-sequence alignments and phylogenetic trees. We performed these analyses on 12,764 gene families from three datasets—24 budding yeasts (1,668 SC-OGs and 1,392 SNAP-OGs) (A), 36 filamentous fungi (4,393 SC-OGs and 2,035 SNAP-OGs) (B), and 26 mammals (321 SC-OGs and 1,775 SNAP-OGs) (C). Principal component analysis revealed striking similarities between SC-OGs and SNAP-OGs in all three datasets. For example, the centroid (i.e., the mean across all metrics and genes) for SC-OGs and SNAP-OGs, which is depicted as an opaque and larger dot, are very close to one another in reduced dimensional space. Supporting this observation, multi-factor analysis of variance with interaction effects of the 6,630 SNAP-OGs and 6,634 SC-OGs revealed no difference between SC-OGs and SNAP-OGs (p = 0.63, F = 0.23, df = 1) and no interaction between the nine properties and SC-OGs and SNAP-OGs (p = 0.16, F = 1.46, df = 8). Multi-factor analysis of variance using an additive model yielded similar results wherein SC-OGs and SNAP-OGs do not differ (p = 0.65, F = 0.21, df = 1). There are also very few outliers of individual SC-OGs and SNAP-OGs, which are represented as translucent dots, in all three panels. For example, SNAP-OGs outliers at the top of panel C are driven by high treeness/RCV values, which is associated with a high signal-to-noise ratio and/or low composition bias [23]; SNAP-OG outliers at the right of panel C are driven by high average bootstrap support values, which is associated with greater tree certainty [66]; and the single SC-OG outlier observed in the bottom right of panel C is driven by a SC-OG with a high degree of violation of a molecular clock [70], which is associated with lower tree certainty [71]. Multiple-sequence alignment and phylogenetic tree properties used in principal component analysis and abbreviations thereof are as follows: average bootstrap support (ABS), degree of violation of the molecular clock (DVMC), relative composition variability, Robinson-Foulds distance (RF distance), alignment length (Aln. len.), the number of parsimony informative sites (PI sites), saturation, treeness (tness), and treeness/RCV (tness/RCV).

We next aimed to determine if SC-OGs and SNAP-OGs have greater phylogenetic information content than a random null expectation. Groups of genes reflecting a random null expectation were constructed by randomly selecting a single sequence from representative species in multi-copy orthologous genes (hereafter referred to as Random-GGs for random combinations of orthologous and paralogous groups of genes) in the budding yeast (N=647), filamentous fungi (N=999), and mammalian (N=954) datasets. Random-GGs were aligned, trimmed, and phylogenetic trees were inferred from the resulting multiple sequence alignments. Random-GG phylogenetic information was also calculated. Across each dataset, significant differences were observed among SC-OGs, SNAP-OGs, and Random-GGs (p < 0.001, F = 189.92, df = 4; Multi-factor ANOVA). Further examination of differences revealed Random-GGs are significantly different compared to SC-OGs and SNAP-OGs (p < 0.001 for both comparisons; Tukey honest significant differences (THSD) test) in the budding yeast dataset. In contrast, SC-OGs and SNAP-OGs are not significantly different (p = 0.42; THSD). The same was also true for the dataset of filamentous fungi and mammals—specifically, Random-GGs were significantly different from SC-OGs and SNAP-OGs (p < 0.001 for each comparison in each dataset; THSD), whereas SC-OGs and SNAP-OGs were not significantly different (p = 1.00 for filamentous fungi dataset; p = 0.42 for dataset of mammals; THSD). Principal component analysis revealed Robinson-Foulds distances (a measure of tree accuracy wherein lower values represent greater tree accuracy), and relative composition variability (a measure of alignment composition bias wherein lower values represent less compositional bias), often drove differences among Random-GGs, SC-OGs, and SNAP-OGs across the datasets. In all datasets, SC-OGs and SNAP-OGs outperformed the null expectation in tree accuracy and were less compositionally biased (Table 1). These findings suggest SNAP-OGs and SC-OGs are similar in phylogenetic information content and outperform the null expectation.

**Table 1.** SC-OGs and SNAP-OGs are more accurate and have less compositional biases than Random-GGs. The first column is the dataset being examined. The second column describes the type of group of genes. The third column is the Robinson-Foulds distances, a measure of tree distance wherein higher values reflect greater inaccuracies. The fourth column is the relative composition variability, a measure of alignment composition bias wherein higher values indicate greater biases. In all datasets, SC-OGs and SNAP-OGs had better scores compared to a null expectation. RF: Robinson-Foulds distance; RCV: relative composition variability. Values represent mean and standard deviations.

| Dataset | OG type | RF distance | RCV |
| --- | --- | --- | --- |
| Budding yeasts | SC-OGs | $0.19 \pm 0.12$ | $0.19 \pm 0.05$ |
| | SNAP-OGs | $0.18 \pm 0.11$ | $0.18 \pm 0.06$ |
|  | Random-GGs | **$0.65 \pm 0.27$** | **$0.27 \pm 0.13$** |
| Filamentous Fungi | SC-OGs | $0.27 \pm 0.13$ | $0.12 \pm 0.05$ |
| | SNAP-OGs | $0.27 \pm 0.12$ | $0.12 \pm 0.06$ |
|  | Random-GGs | **$0.87 \pm 0.11$** | **$0.21 \pm 0.13$** |
| Mammals | SC-OGs | $0.56 \pm 0.22$ | $0.13 \pm 0.06$ |
| | SNAP-OGs | $0.51 \pm 0.23$ | $0.11 \pm 0.07$ |
|  | Random-GGs | **$0.61 \pm 0.30$** | **$0.15 \pm 0.10$** |

### SC-OGs and SNAP-OGs have similar performances in complex datasets

Complex biological processes and datasets pose a serious challenge for identifying markers for molecular evolution studies. To test the efficacy of OrthoSNAP in scenarios of complex evolutionary histories and datasets, we executed the same workflow described above—ortholog calling, sequence alignment, trimming, tree inference, and SNAP-OG detection—on three new datasets: (1) 30 plants, which are known to complex histories of gene duplication and loss [24–26]; (2) 30 budding yeast species wherein half of the species originated from a hybridization event that gave rise to a whole genome duplication followed by complex patterns of loss and duplication [27–30]; and (3) 20 choanoflagellate transcriptomes, which contain thousands more transcripts than genes [31,32]; for orthology inference software, multiple transcripts per gene appear similar to artificial gene duplicates.

Corroborating previous results, OrthoSNAP successfully identified SNAP-OGs that can be used downstream for molecular evolution analyses. Specifically, using a species-occupancy threshold of 50% in the plant, budding yeast, and choanoflagellate datasets, 653, 1,334, and 2,087 SNAP-OGs were identified, respectively (Table 2). In comparison, 15 SC-OGs were identified in the plant dataset; 2,782 in the budding yeast dataset; and 390 in the choanoflagellate dataset. To explore the impact of orthogroup occupancy, SNAP-OGs were also identified using a minimum occupancy threshold of four taxa. This resulted in the identification of substantially more SNAP-OGs: 15,854 in plants; 4,199 in budding yeasts; and 11,556 in choanoflagellates. Furthermore, these were substantially higher than the number of SC-OGs identified using a minimum orthogroup occupancy of four taxa: 200 in plants; 3,566 in budding yeasts; and 2,438 in choanoflagellates. These findings support previous observations that incorporating OrthoSNAP into ortholog identification workflows can substantially increase the number of available loci.

**Table 2.** OrthoSNAP identifies SNAP-OGs in complex datasets. SC-OG identification can be difficult due to complex evolutionary histories (e.g., hybridization, whole genome duplication, and complex patterns of gene duplication and loss such as in the datasets of budding yeasts and plants) and analytical artifacts (e.g., transcriptomes with more transcripts than genes such as the choanoflagellate dataset). OrthoSNAP successfully identified SNAP-OGs in each dataset. Lowering the occupancy threshold of a SNAP-OG to a minimum of four enabled the identification of substantially more SNAP-OGs.

| **Dataset** | **Challenge** | **Total**<br>**OGs** | **SC-OGs**<br>**(50% min.**<br>**occupancy**<br>**threshold)** | **SNAP-**<br>**OGs (50%**<br>**min.**<br>**occupancy**<br>**threshold)** | **SC-OGs**<br>**(4 species**<br>**min.**<br>**threshold)** | **SNAP-**<br>**OGs (4**<br>**species**<br>**min.**<br>**threshold)** |
| --- | --- | --- | --- | --- | --- | --- |
| Plants (N=30) | Evolutionary histories with extensive gene duplication and loss events | 83,034 | 15 | 653 | 200 | 15,854 |
| Budding yeasts (N=30) | Half of the species used experienced hybridization and whole-genome duplication followed by extensive loss of paralogs | 11,422 | 2,782 | 1,334 | 3,566 | 4,199 |
| Choanoflagellates (N=20) | Transcriptomes, where often multiple transcripts correspond to a single protein- coding gene | 274,028 | 390 | 2,087 | 2,438 | 11,556 |

### SC-OGs and SNAP-OGs have similar patterns of support in a contentious branch in the tree of life

To further evaluate the information content of SNAP-OGs, we compared patterns of support among SC-OGs and SNAP-OGs in a difficult-to-resolve branch in the tree of life. Specifically, we evaluated the support between three hypotheses concerning deep evolutionary relationships among eutherian mammals: (1) Xenarthra (eutherian mammals from the Americas) and Afrotheria (eutherian mammals from Africa) are sister to all other Eutheria [33,34]; (2) Afrotheria are sister to all other Eutheria [35,36]; and (3) Xenarthra are sister to a clade of both Afrotheria and Eutheria (Fig 4A). Resolution of this conflict has important implications for understanding the historical biogeography of these organisms. To do so, we first obtained protein-coding gene sequences from six Afrotheria, two Xenarthra, 12 other Eutheria, and eight outgroup taxa from NCBI (Table S7), which represent all annotated and publicly genome assemblies at the time of this study (Table S8). Using the protein translations of these gene sequences as input to OrthoFinder, we identified 252 SC-OGs shared across taxa; application of OrthoSNAP identified an additional 1,428 SNAP-OGs, which represents a greater than five-fold increase in the number of gene markers for this dataset (Table S8). There was variation in the number of SNAP-OGs identified per orthologous group of genes (S6 Fig). The highest number of SNAP-OGs identified in an orthologous group of genes was 10, which was a gene family of olfactory receptors; olfactory receptors are known to have expanded in the evolutionary history of eutherian mammals [8]. The best fitting substitution models were similar between SC-OGs and SNAP-OGs (S7 Fig).

**Figure 4.**
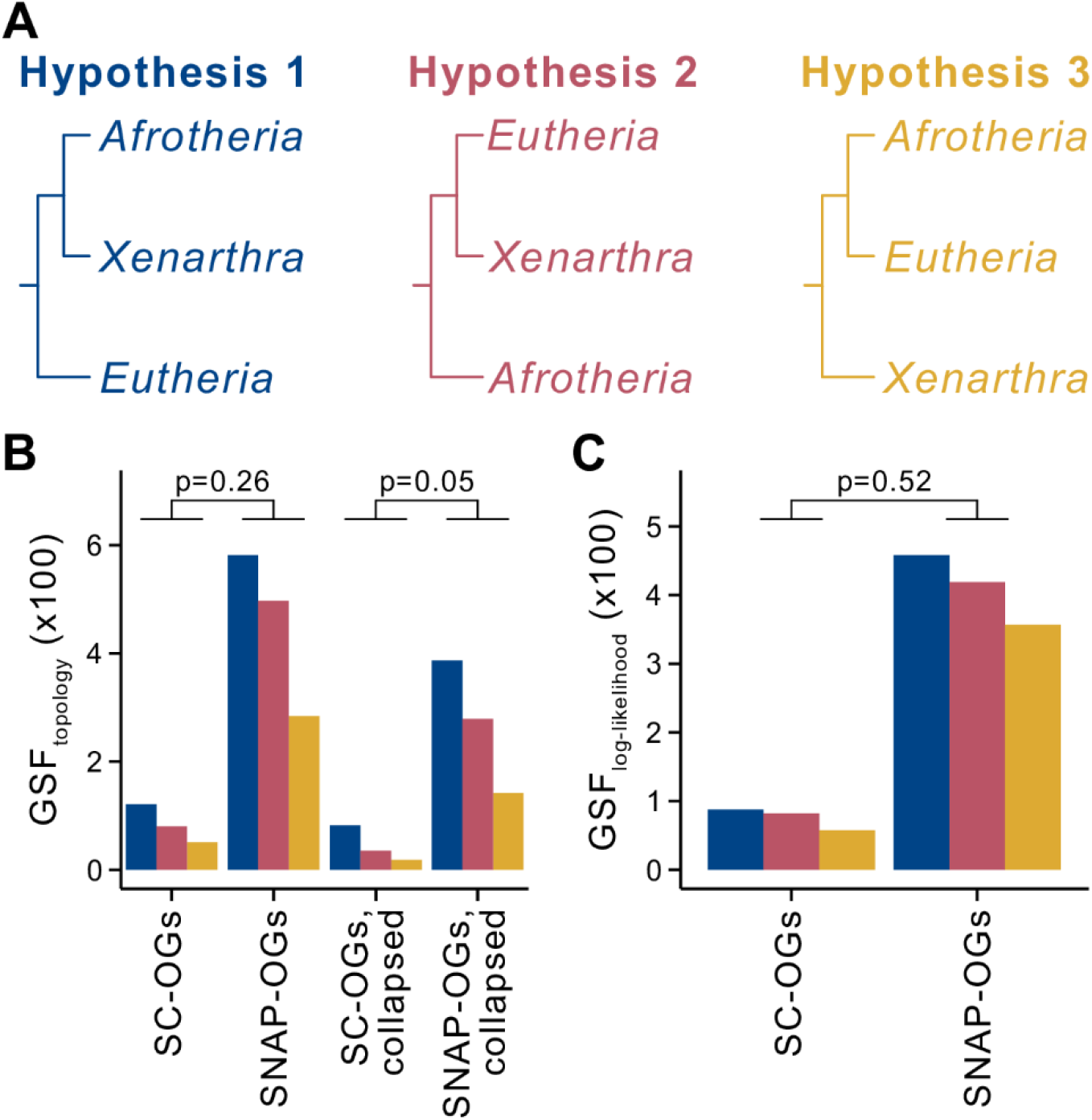
SC-OGs and SNAP-OGs display similar patterns of support in a contentious branch concerning deep evolutionary relationships among eutherian mammals. (A) Two leading hypotheses for the evolutionary relationships among Eutheria, which have implications for the evolution and biogeography of the clade, are that Afrotheria and Xenarthra are sister to all other Eutheria (hypothesis one; blue) and that Afrotheria are sister to all other Eutheria (hypothesis two; pink). The third possible, but less well supported topology, is that Xenarthra are sister to Eutheria and Afrotheria. (B) Comparison of gene support frequency (GSF) values for the three hypotheses among 252 SC-OGs and 1,428 SNAP-OGs using an α level of 0.01 revealed no differences in support (p = 0.26, Fisher’s exact test with Benjamini-Hochberg multi-test correction). Comparison after accounting for gene tree uncertainty by collapsing bipartitions with ultrafast bootstrap approximation support lower than 75 (SC-OGs collapsed vs. SNAP-OGs collapsed) also revealed no differences (p = 0.05; Fisher’s exact test with Benjamini-Hochberg multi-test correction). (C) Examination of the distribution of frequency of topology support using gene-wise log-likelihood scores revealed no difference between SNAP-OGs and SC-OGs support for the three topologies (p = 0.52; Fisher’s exact test).

Two independent tests examining support between alternative hypotheses of deep evolutionary relationships among eutherian mammals revealed similar patterns of support between SC-OGs and SNAP-OGs. More specifically, no differences were observed in gene support frequencies—the number of genes that support one of three possible hypotheses at a given branch in a phylogeny—without or with accounting for single-gene tree uncertainty by collapsing branches with low support values (p = 0.26 and p = 0.05, respectively; Fisher’s exact test with Benjamini-Hochberg multi-test correction; Fig 4B; Table S9). A second test of single-gene support was conducted wherein individual gene log likelihoods were calculated for each of the three possible topologies. The frequency of gene-wise support for each topology was determined. No differences were observed in gene support frequency using the log likelihood approach (p = 0.52, respectively; Fisher’s exact test). Examination of patterns of support in a contentious branch in the tree of life using two independent tests revealed SC-OGs and SNAP-OGs are similar and further supports the observation that they contain similar phylogenetic information.

In summary, 415,129 orthologous groups of genes across seven eukaryotic datasets contained 9,821 SC-OGs; application of OrthoSNAP identified an additional 10,704 SNAP-OGs, thereby more than doubling the number of gene markers. Comprehensive comparison of the phylogenetic information content among SC-OGs and SNAP-OGs revealed no differences in phylogenetic information content. Strikingly, this observation held true across datasets with complex evolutionary histories and when conducting hypothesis testing in a difficult-to-resolve branch in the tree of life. These findings suggest that SNAP-OGs may be useful for diverse studies of molecular evolution ranging from genome-wide surveys of selection, phylogenomic investigations, gene-gene coevolution analyses, and others.

## Discussion

Molecular evolution studies typically rely on SC-OGs. Recently developed methods can integrate gene families of orthologs and paralogs into species tree inference but are not designed to broadly facilitate the retrieval of gene markers for molecular evolution analyses. Furthermore, the phylogenetic information content of gene families of orthologs and paralogs remains unknown. This observation underscores the need for algorithms that can identify SC-OGs nested within larger gene families, which can be in turn be incorporated into diverse molecular evolution analyses, and a comprehensive assessment of their phylogenetic properties.

To address this need, we developed OrthoSNAP, a tree splitting and pruning algorithm that identifies SNAP-OGs, which refers to SC-OGs nested within larger gene families wherein species specific inparalogs have also been pruned. Comprehensive examination of the phylogenetic information content of SNAP-OGs and SC-OGs from seven empirical datasets of diverse eukaryotic species revealed that their content is similar. Inclusion of SNAP-OGs increased the size of all seven datasets, sometimes substantially. We note that our results are qualitatively similar to those reported recently by Smith et al. [37], which retrieved SC-OGs nested within larger families from 26 primates and examined their performance in gene tree and species tree inference. Three noteworthy differences are that we also conduct species-specific inparalog trimming, provide a user-friendly command-line software for SNAP-OG identification, and evaluated the phylogenetic information content of SNAP-OGs and SC-OGs across seven diverse phylogenomic datasets. We also note that our algorithm can account for diverse types of paralogy—outparalogs, inparalogs, and species-specific inparalogs—whereas other software like PhyloTreePruner, which only conducts species-specific inparalog trimming [19], and Agalma, which identifies single-copy outparalogs and inparalogs [20], can account for some, but not all, types of paralogs (Table S10). Another difference between OrthoSNAP and other approaches is that Agalma and PhyloTreePruner both require rooted phylogenies. In contrast, OrthoSNAP will automatically midpoint root phylogenies or accept pre-rooted phylogenies as input. Furthermore, these algorithms are not designed to handle transcriptomic data wherein multiple transcripts per gene will be interpreted as multi-copy orthologs. Thus, OrthoSNAP allows for greater user flexibility and accounts for more diverse scenarios, leading to, at least in some instances, the identification of more loci for downstream analyses (S8 Fig). Notably, these software are also different from sequence similarity graph-based inferences of subgroups of single-copy orthologous genes—such as the algorithm implemented in OMA [21]. Finally, our results, together with other studies, demonstrate the utility of SC-OGs that are nested within larger families [15,20,37,38].

Despite the ability of OrthoSNAP to identify additional loci for molecular evolution analyses, there were instances wherein SNAP-OGs were not identified in multi-copy orthologous groups of genes. We discuss three reasons that contribute to why SNAP-OGs could not be identified among some genes—specifically, gene families with sequence data from <50% of the taxa; gene families with complex evolutionary histories (for example, HGT and duplication/loss patterns); and gene families with evolutionary histories that differ from the species tree (for example, due to analytical factors, such as sampling and systematic error, or biological factors, such as lineage sorting or introgression). Notably, the first reason can, but does not always, result in inability to infer SNAP-OGs and can be, to a certain extent, addressed (e.g., by lowering the orthogroup occupancy threshold in OrthoSNAP), whereas the other two reasons are more challenging because they often result in a genuine absence of single-copy orthologs. Furthermore, the actual number of single-copy orthologs (either those nested within multi-copy orthologs or not) for any given group of organisms is not known, making it difficult to determine how many SNAP-OGs and SC-OGs one should expect to recover. Notably, this issue has long challenged researchers, even when ortholog identification is performed by also taking genome synteny into account [27].

Next, we discuss some practical considerations when using OrthoSNAP. In the present study, we inferred orthology information using OrthoFinder [39], but several other approaches can be used upstream of OrthoSNAP. For example, other graph-based algorithms such as OrthoMCL and OMA [21,40] or sequence similarity-based algorithms such as orthofisher [41], can be used to infer gene families. Similarly, sequence similarity search algorithms like BLAST+ [42], USEARCH [43], and HMMER [44], can be used to retrieve homologous sets of sequences that are used as input for OrthoSNAP. Other considerations should also be taken during the multi-copy tree inference step. For example, inferring phylogenies for all orthologous groups of genes may be a computationally expensive task. Rapid tree inference software—such as FastTree or IQTREE with the “-fast” parameter [45,46]—may expedite these steps (but users should be aware that this may result in a loss of accuracy in inference [47]).

We suggest employing “best practices” when inferring groups of putatively orthologous genes, including SNAP-OGs. Specifically, orthology information can be further scrutinized using phylogenetic methods. Orthology inference errors may occur upstream of OrthoSNAP; for example, SNAP-OGs may be susceptible to erroneous inference of orthology during upstream clustering of putatively orthologous genes. One method to identify putatively spurious orthology inference is by identifying long terminal branches [48]. Terminal branches of outlier length can be identified using the “spurious_sequence” function in PhyKIT [49]. Other tools, such as PhyloFisher, UPhO, and other orthology inference pipelines employ similar strategies to refine orthology inference [50–52]. Lastly, we acknowledge that future iterations of OrthoSNAP may benefit from incorporating additional layers of information, such as sequence similarity scores or synteny. Even though OrthoSNAP did identify SNAP-OGs in some complex datasets where synteny has previously been very helpful, such as the budding yeast dataset, other ancient and rapidly evolving lineages may benefit from synteny analysis to dissect complex relationships of orthology [48,53–55].

Taken together, we suggest that OrthoSNAP is useful for retrieving single-copy orthologous groups of genes from gene family data and that the identified SNAP-OGs have similar phylogenetic information content compared to SC-OGs. In combination with other phylogenomic toolkits, OrthoSNAP may be helpful for reconstructing the tree of life and expanding our understanding of the tempo and mode of evolution therein.

## Methods

### OrthoSNAP availability and documentation

OrthoSNAP is available under the MIT license from GitHub (https://github.com/JLSteenwyk/orthosnap), PyPi (https://pypi.org/project/orthosnap), and the Anaconda cloud (https://anaconda.org/JLSteenwyk/orthosnap). OrthoSNAP is also freely available to use via the LatchBio (https://latch.bio/) cloud-based console (dedicated interface link: https://console.latch.bio/explore/65606/info). Documentation describes the OrthoSNAP algorithm, parameters, and provides user tutorials (https://jlsteenwyk.com/orthosnap).

### OrthoSNAP algorithm description and usage

We next describe how OrthoSNAP identifies SNAP-OGs. OrthoSNAP requires two files as input: one is a FASTA file that contains two or more homologous sequences in one or more species and the other the corresponding gene family phylogeny in Newick format. In both the FASTA and Newick file, users must follow a naming scheme—wherein species, strain, or organism identifiers and gene sequences identifiers are separated by a vertical bar (also known as a pipe character or “|”)—which allows OrthoSNAP to determine which sequences were encoded in the genome of each species, strain, or organism. After initiating OrthoSNAP, the gene family phylogeny is first mid-point rooted (unless the user specifies the inputted phylogeny is already rooted) and then SNAP-OGs are identified using a tree-traversal algorithm. To do so, OrthoSNAP will loop through the internal branches in the gene family phylogeny and evaluate the number of distinct taxa identifiers among children terminal branches. If the number of unique taxon identifiers is greater than or equal to the orthogroup occupancy threshold (default: 50% of total taxa in the inputted phylogeny; users can specify an integer threshold), then all children branches and termini are examined further; otherwise, OrthoSNAP will examine the next internal branch. Next, OrthoSNAP will collapse branches with low support (default: 80, which is motivated by using ultrafast bootstrap approximations [56] to evaluate bipartition support; users can specify an integer threshold) and conduct species-specific inparalog trimming wherein the longest sequence is maintained, a practice common in transcriptomics. However, users can specify whether the shortest sequence or the median sequence (in the case of three or more sequences) should be kept instead. Users can also pick which species-specific inparalog to keep based on branch lengths (the longest, shortest, or median branch length in the case of having three or more sequences). Species-specific inparalogs are defined as sequences encoded in the same genome that are sister to one another or belong to the same polytomy [19]. The resulting set of sequences is examined to determine if one species, strain, or organism is represented by one sequence and ensure these sequences have not yet been assigned to a SNAP-OG. If so, they are considered a SNAP-OG; if not, OrthoSNAP will examine the next internal branch. When SNAP-OGs are identified, FASTA files of SNAP-OG sequences are outputted. Users can also output the subtree of the SNAP-OG using an additional argument.

The principles of the OrthoSNAP algorithm are also described using the following pseudocode: FOR internal branch in midpoint rooted gene family phylogeny:

> IF orthogroup occupancy among children termini is greater than or equal to orthogroup occupancy threshold;

>> Collapse poorly supported bipartitions and trim species-specific inparalogs;

>> IF each species, strain, or organism among the trimmed set of species, strains, or organisms is represented by only one sequence and no sequences being examined have been assigned to a SNAP-OG yet;

>>> Sequences represent a SNAP-OG and are outputted to a FASTA file

>> ELSE

>>> examine next internal branch

> ELSE

>> examine next internal branch

ENDFOR

To enhance the user experience, arguments or default values are printed to the standard output, a progress bar informs the user of how of the analysis has been completed, and the number of SNAP-OGs identified as well as the names of the outputted FASTA files are printed to the standard output.

### Development practices and design principles to ensure long-term software stability

Archival instabilities among software threatens the reproducibility of bioinformatics research [57]. To ensure long-term stability of OrthoSNAP, we implemented previously established rigorous development practices and design principles [41,49,58,59]. For example, OrthoSNAP features a refactored codebase, which facilitates debugging, testing, and future development. We also implemented a continuous integration pipeline to automatically build, package, and install OrthoSNAP across Python versions 3.7, 3.8, and 3.9. The continuous integration pipeline also conducts 57 unit and integration tests, which span 95.90% of the codebase and ensure faithful function of OrthoSNAP.

### Dataset generation

To generate a dataset for identifying SNAP-OGs and comparing them to SC-OGs, we first identified putative groups of orthologous genes across four empirical datasets. To do so, we first downloaded proteomes for each dataset, which were obtained from publicly available repositories on NCBI (Table S1 and S7) or figshare [48]. Each dataset varied in its sampling of sequence diversity and in the evolutionary divergence of the sampled taxa. The dataset of 24 budding yeasts spans approximately 275 million years of evolution [48]; the dataset of 36 filamentous fungi spans approximately 94 million years of evolution [60]; the dataset of 26 mammals spans approximately 160 million years of evolution [61]; and the dataset of 28 eutherian mammals—which was used to study the contentious deep evolutionary relationships among eutherian mammals—concerns an ancient divergence that occurred approximately 160 million years ago [62]. Putatively orthologous groups of genes were identified using OrthoFinder, v2.3.8 [39], with default parameters, which resulted in 46,645 orthologous groups of genes with at least 50% orthogroup occupancy (Table S8).

To infer the evolutionary history of each orthologous group of genes, we first individually aligned and trimmed each group of sequences using MAFFT, v7.402 [63], with the “auto” parameter and ClipKIT, v1.1.3 [58], with the “smart-gap” parameter, respectively. Thereafter, we inferred the best-fitting substitution model using Bayesian information criterion and evolutionary history of each orthologous group of genes using IQ-TREE2, v2.0.6 [46]. Bipartition support was examined using 1,000 ultrafast bootstrap approximations [56].

To identify SNAP-OGs, the FASTA file and associated phylogenetic tree for each gene family with multiple homologs in one or more species was used as input for OrthoSNAP, v0.0.1 (this study). Across 40,011 gene families with multiple homologs in one or more species in all datasets, we identified 6,630 SNAP-OGs with at least 50% orthogroup occupancy (S1 Fig; Table S8). Unaligned sequences of SNAP-OGs were then individually aligned and trimmed using the same strategy as described above. To determine gene families that were SC-OGs, we identified orthologous groups of genes with at least 50% orthogroup occupancy and each species, strain, or organism was represented by only one sequence— 6,634 orthologous groups of genes were SC-OGs.

### Measuring and comparing information content among SC-OGs and SNAP-OGs

To compare the information content of SC-OGs and SNAP-OGs, we calculated nine properties of multiple sequence alignments and phylogenetic trees associated with robust phylogenetic signal in the budding yeasts, filamentous fungi, and mammalian datasets (Table S4). More specifically, we calculated information content from phylogenetic trees such as measures of tree certainty (average bootstrap support), accuracy (Robinson-Foulds distance (Robinson and Foulds, 1981)), signal-to-noise ratios (treeness (Phillips and Penny, 2003)), and violation of clock-like evolution (degree of violation of a molecular clock or DVMC (Liu et al., 2017)). Information content was also measured among multiple sequence alignments by examining alignment length and the number of parsimony-informative sites, which are associated with robust and accurate inferences of evolutionary histories (Shen et al., 2016) as well as biases in sequence composition (RCV (Phillips and Penny, 2003)). Lastly, information content was also evaluated using metrics that consider characteristics of phylogenetic trees and multiple sequence alignments such as the degree of saturation, which refers to multiple substitutions in multiple sequence alignments that underestimate the distance between two taxa (Philippe et al., 2011), and treeness / RCV, a measure of signal-to-noise ratios in phylogenetic trees and sequence composition biases (Phillips and Penny, 2003). For tree accuracy, phylogenetic trees were compared to species trees reported in previous studies [48,60,61]. All properties were calculated using functions in PhyKIT, v1.1.2 [49]. The function used to calculate each metric and additional information are described in Table S4.

Principal component analysis across the nine properties that summarize phylogenetic information content was used to qualitatively compare SC-OGs and SNAP-OGs in reduced dimensional space. Principal component analysis, visualization, and determination of property contribution to each principal component was conducted using factoextra, v1.0.7 [64], and FactoMineR, v2.4 [65], in the R, v4.0.2 (https://cran.r-project.org/), programming environment. Statistical analysis using a multi-factor ANOVA was used to quantitatively compare SC-OGs and SNAP-OGs using the res.aov() function in R.

Information theory-based approaches were used to evaluate incongruence among SC-OGs and SNAP-OGs phylogenetic trees. More specifically, we calculated tree certainty and tree certainty-all [66–68], which are conceptually similar to entropy values and are derived from examining support among a set of gene trees and the two most supported topologies or all topologies that occur with a frequency of ≥5%, respectively. More simply, tree certainty values range from 0 to 1 in which low values are indicative of low congruence among gene trees and high values are indicative of high congruence among gene trees. Tree certainty and tree certainty-all values were calculated using RAxML, v8.2.10 [69].

To examine patterns of support in a contentious branch concerning deep evolutionary relationships among eutherian mammals, we calculated gene support frequencies and ΔGLS. Gene support frequencies were calculated using the “polytomy_test” function in PhyKIT, v1.1.2 [49]. To account for uncertainty in gene tree topology, we also examined patterns of gene support frequencies after collapsing bipartitions with ultrafast bootstrap approximation support lower than 75 using the “collapse” function in PhyKIT. To calculate gene-wise log likelihood values, partition log-likelihoods were calculated using the “wpl” parameter in IQ-TREE2 [46], which required as input a phylogeny in Newick format that represented either hypothesis one, two, or three (Fig 4A) and a concatenated alignment of SC-OGs and SNAP-OGs with partition information. Thereafter, the log likelihood values were used to assign genes to the topology they best supported. Inconclusive genes, defined as having a gene-wise log likelihood difference of less than 0.01, were removed.

The same methodologies—orthology inference, multiple-sequence alignment, trimming, tree inference, SNAP-OG identification, and phylogenetic information content calculations—were also applied to three additional datasets that represent complex datasets. Specifically, 30 plants (with a history of extensive gene duplication and loss events), 30 budding yeast species (15 of which experienced whole-genome duplication), and 20 choanoflagellate transcriptomes (where typically multiple transcripts correspond to a single protein-coding gene) [31,32].

## Data Availability

All results and data presented in this study are available from figshare (doi: 10.6084/m9.figshare.16875904).

## Acknowledgements

We thank the Rokas lab for helpful discussion and feedback. J.L.S. and A.R. were funded by the Howard Hughes Medical Institute through the James H. Gilliam Fellowships for Advanced Study program. Research in A.R.’s lab is supported by grants from the National Science Foundation (DEB-2110404), the National Institutes of Health/National Institute of Allergy and Infectious Diseases (R56 AI146096 and R01 AI153356), and the Burroughs Wellcome Fund.

