## Supplementary Figures for "OrthoSNAP: a tree splitting and pruning algorithm for retrieving single-copy orthologs from gene family trees"

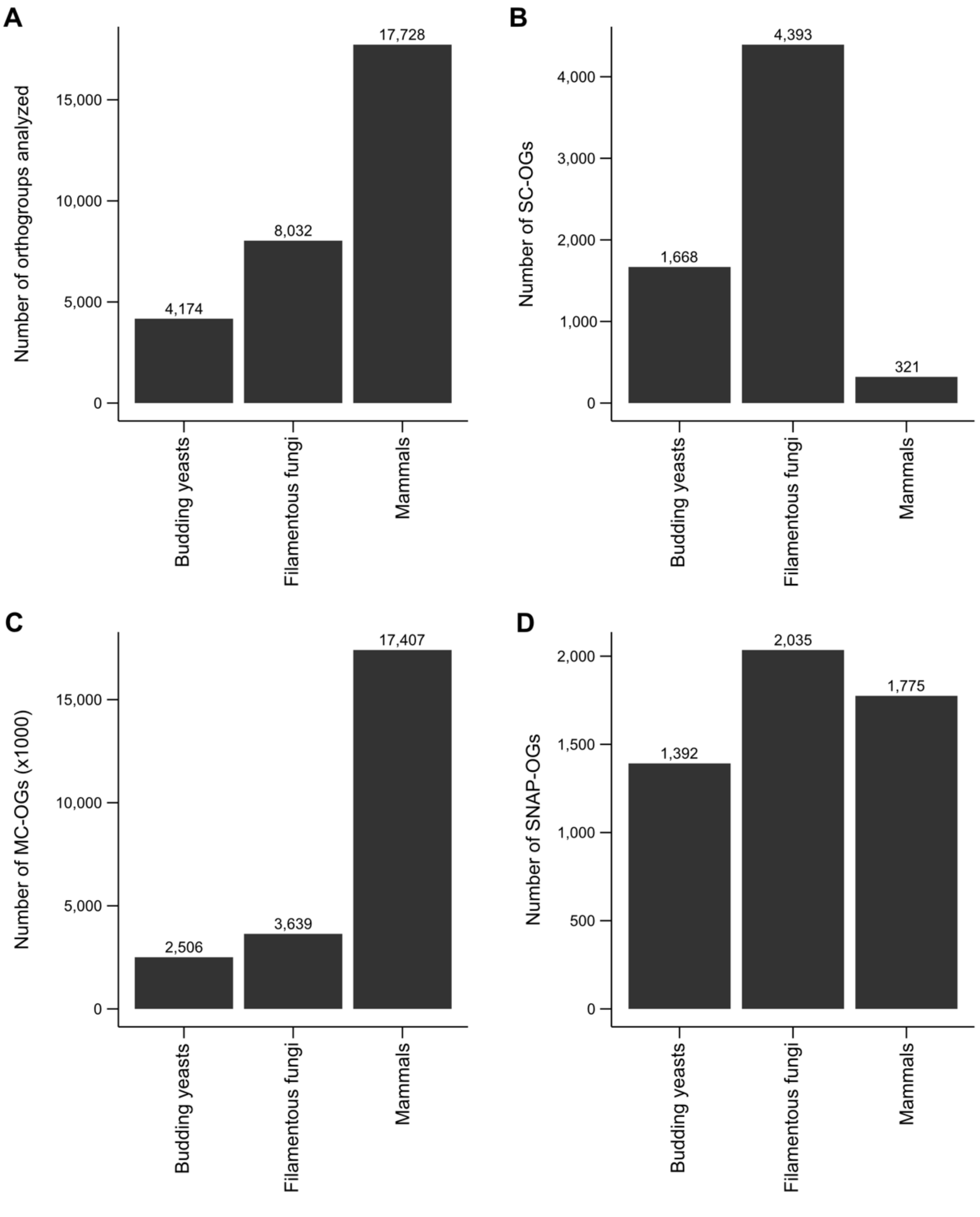


**Supplementary figure 1. Numbers of orthogroups, single-copy orthogroups, orthogroups with one or more homologs in one species, and the number of SNAP-OGs identified for each dataset.** (A) The total number of orthogroups with at least 50% ortholog occupancy for each dataset. (B) The number of single-copy orthologs (SC-OGs) for each dataset (with at least 50% taxon occupancy). (C) The number of multi-copy orthologs (or orthologous groups of genes wherein one or more species is represented by two or more sequences; MC-OGs) for each dataset (with at least 50% taxon occupancy). (D) The number of

SNAP-OGs identified in each dataset (with at least 50% taxon occupancy). Note that the numbers depicted in panel A reflect the sum of the numbers of SC-OGs and MC-OGs in panels B and C.

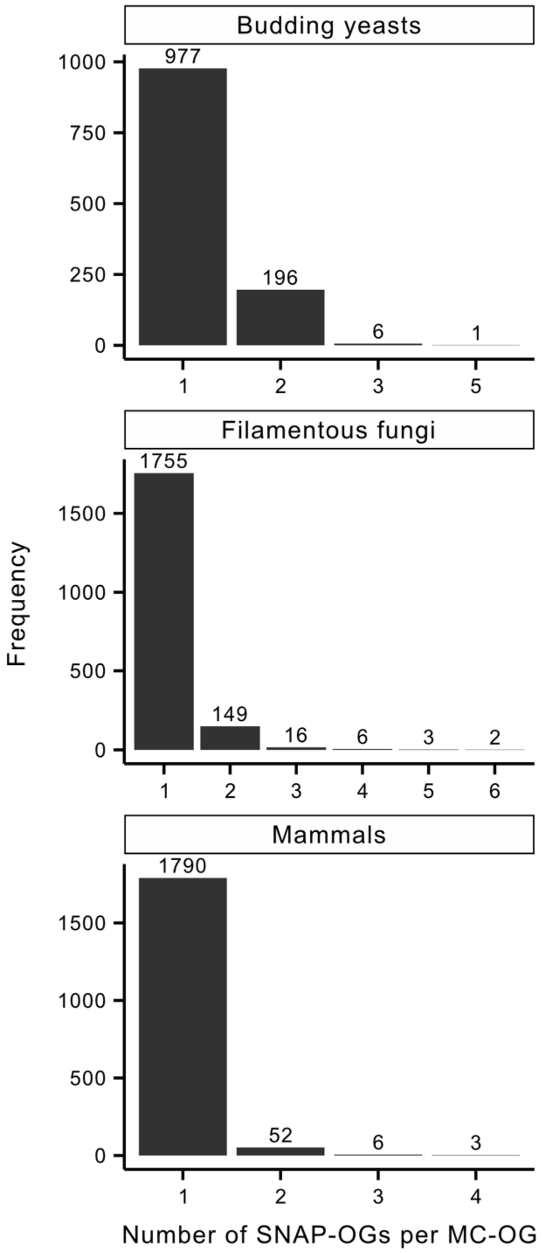


**Supplementary figure 2. The number of SNAP-OGs identified in orthologous groups of genes with two or more homologs in one or more species.** The number of SNAP-OGs per orthologous group of genes is depicted on the x-axis. For example, in the budding yeasts dataset, 977 gene families had one SNAP-OG each. The highest number of SNAP-OGs identified in a single orthologous group of genes in each dataset were as follows: in budding yeasts, five SNAP-OGs were identified in one orthologous group of genes that encode transcriptional activators; in filamentous fungi, five SNAP-OGs were identified in each of two orthologous groups of genes that encode multi-facilitator superfamily transporters and amino acid permeases; and in mammals, four SNAP-OGs were identified in each of three orthologous groups of genes that encode voltage-gated potassium channels, casein kinases, and a tropomyosin family of actin-binding proteins.

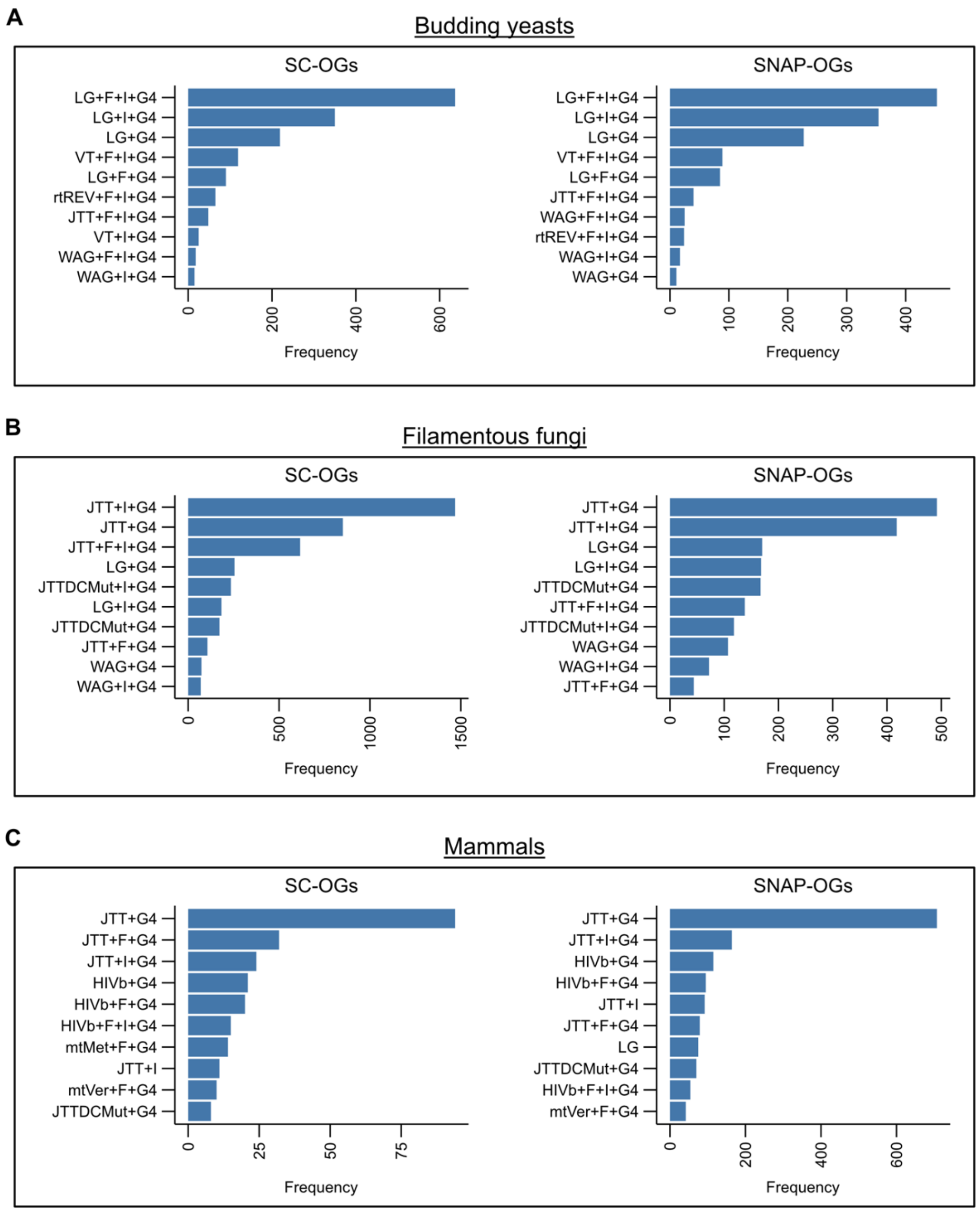


**Supplementary figure 3. The 10 most frequent best-fitting substitutions models are similar between SC-OGs and SNAP-OGs.** The top 10 most frequently observed best-fitting substitutions models were similar between SC-OGs and SNAP-OGs among (A) 1,668 SC-OGs and 1,392 SNAP-OGs in budding yeasts, (B) 4,393 SC-OGs and 2,035 SNAP-OGs in filamentous fungi, and (C) 321 SC-OGs and 1,775 SNAP-OGs in mammals. For example, the LG+F+I+G4 model was the most frequently observed best-fitting substitution model in SC-OGs and SNAP-OGs from budding yeasts.

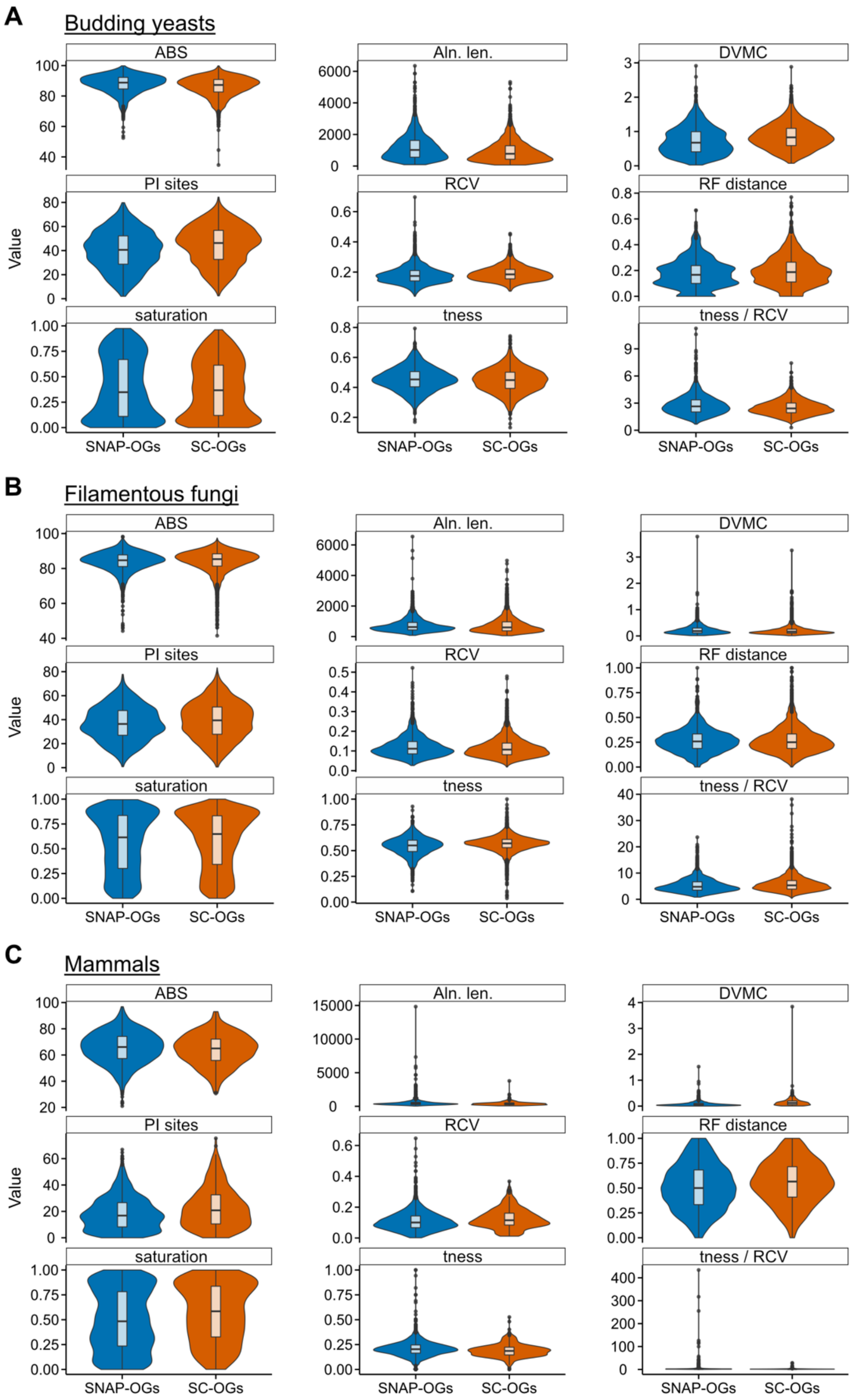


**Supplementary figure 4. Distributions of information content among SNAP-OGs and SC-OGs.**

Boxplot and violin plot distributions of nine properties representative of phylogenetic information are depicted SNAP-OGs (blue) and SC-OGs (orange) in the (A) 1,668 SC-OGs and 1,392 SNAP-OGs in budding yeasts, (B) 4,393 SC-OGs and 2,035 SNAP-OGs in filamentous fungi, and (C) 321 SC-OGs and 1,775 SNAP-OGs in mammals. Abbreviations are as follows: average bootstrap support (ABS), degree of violation of the molecular clock (DVMC), relative composition variability, Robinson-Foulds distance (RF distance), alignment length (Aln. len.), the number of parsimony informative sites (PI sites), saturation, treeness (tness), and treeness/RCV (tness/RCV).

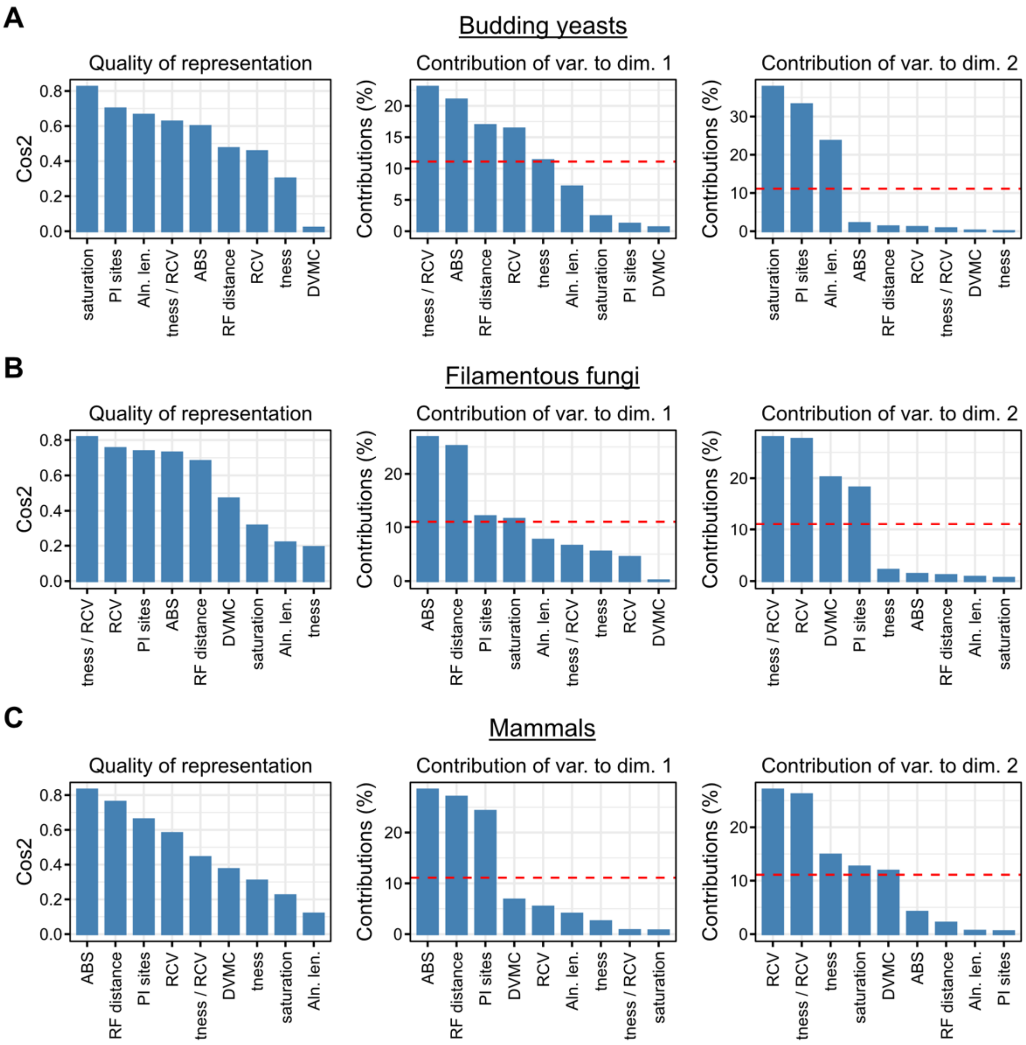


**Supplementary figure 5. Quality of representation and contributions of properties of phylogenetic information content during principal component analysis.** Principal component analysis was used to qualitatively compare the similarities and differences between SNAP-OGs and SC-OGs (Fig 3). The leftmost figure in each panel of budding yeasts (A), filamentous fungi (B), and mammals (C) represents the quality of representation for each property across all principal components. The next two figures depict the contribution of each property (or variable) to the first and second dimension in reduced dimensional space. The red dashed line represents equal contributions from each variable.

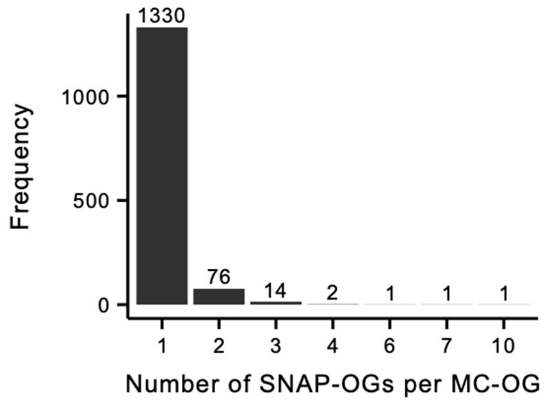


**Supplementary figure 6. The number of SNAP-OGs identified in an orthologous group of genes with two or more homologs in one or more species for the dataset used to examine a contentious branch in the tree of life.** The number of SNAP-OGs per orthologous group of genes is depicted on the x-axis. For example, a single SNAP-OG was identified in 1,330 gene families with two or more homologs in one or more species, whereas four SNAP-OGs were identified in two gene families with two or more homologs in one or more species.

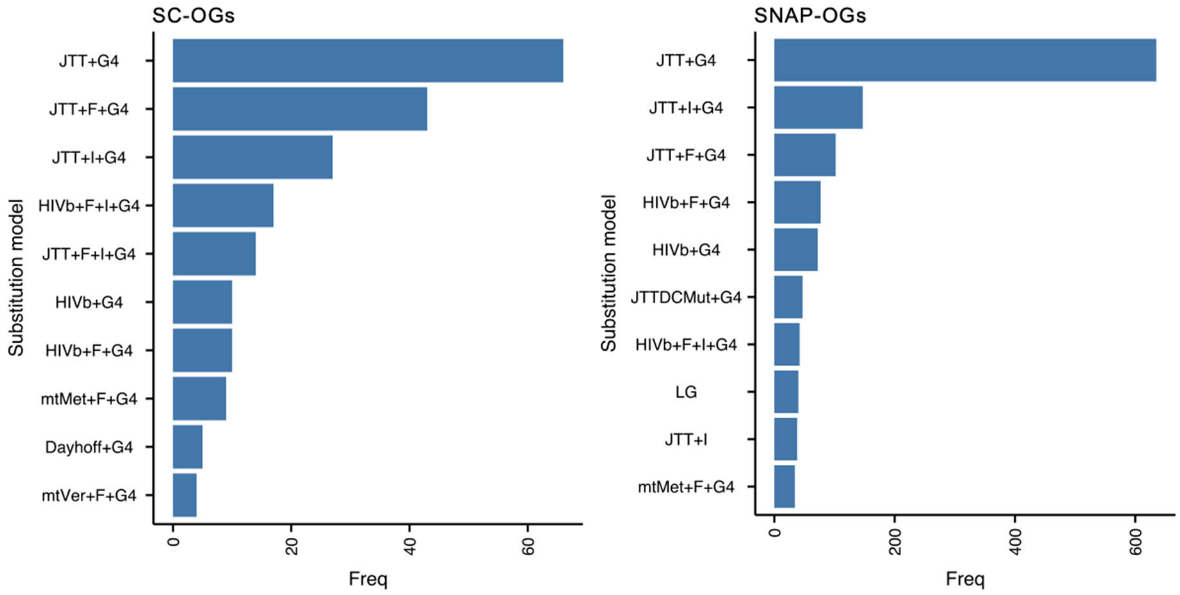


**Supplementary figure 7. The ten most frequently observed best-fitting substitutions models are similar between SC-OGs and SNAP-OGs in the dataset used to examine a contentious branch in the tree of life.** Similar best-fitting substitutions models were observed between 252 SC-OGs and 1,428 SNAP-OGs in a dataset of mammals, which was used to investigate patterns of support in a contentious branch in the tree of life concerning deep evolutionary relationships among placental mammals.

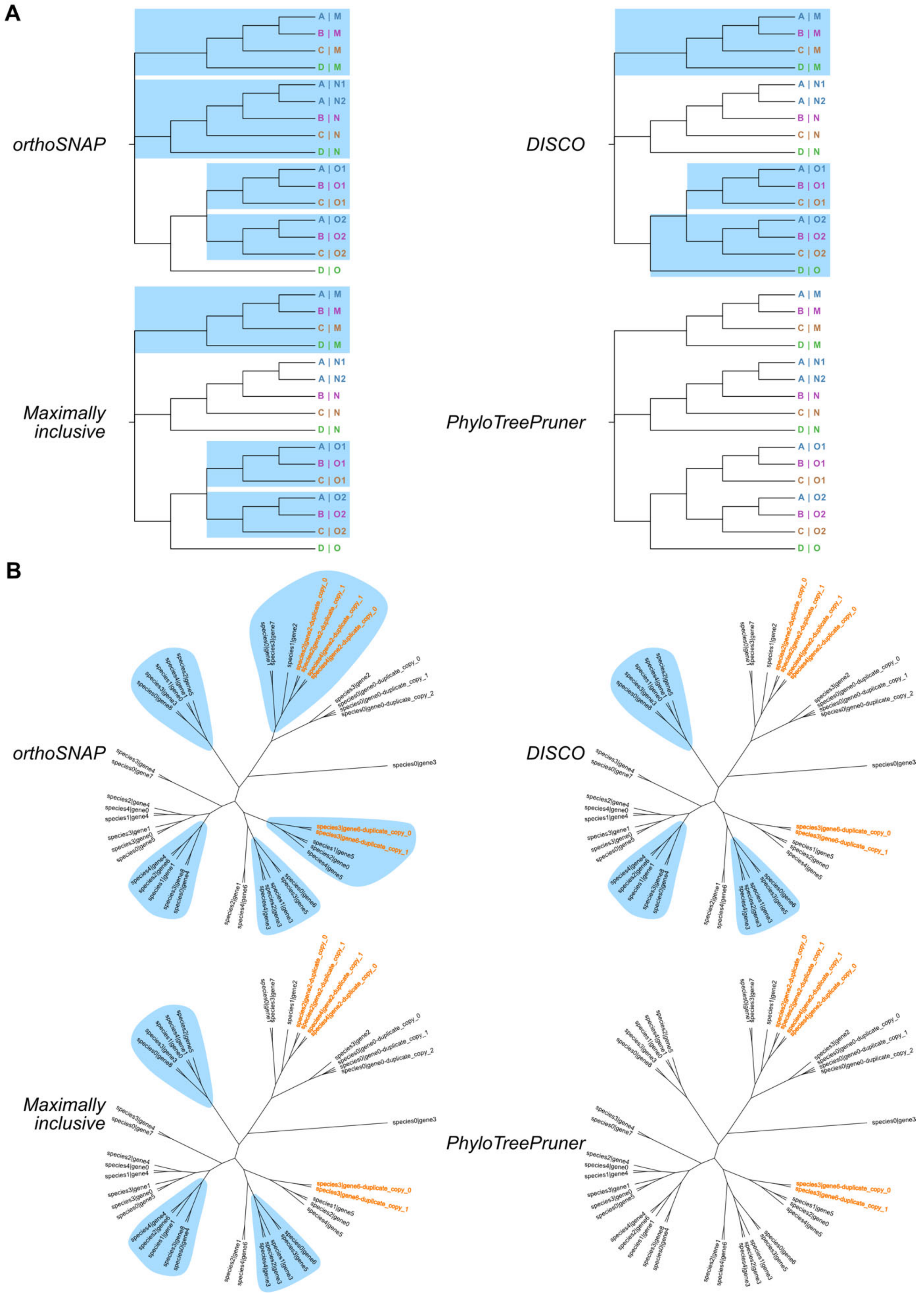
A
orthoSNAP
DISCO
Maximally inclusive
PhyloTreePruner
A | M
B | M
C | M
D | M
A | N1
A | N2
B | N
C | N
D | N
A | O1
B | O1
C | O1
A | O2
B | O2
C | O2
D | O
B
orthoSNAP
DISCO
Maximally inclusive
PhyloTreePruner
species0|gene3

**Supplementary figure 8. Cartoon comparison of different tree decomposition algorithms.** Using the phylogeny presented in Figure 1B (panel A) and Figure 2B (panel B), different tree decomposition algorithms are compared. (A) OrthoSNAP will identify four SNAP-OGs whereas DISCO and the maximally inclusive strategies will each identify three subgroups of orthologous genes. PhyloTreePruner will not identify any subgroups of single-copy orthologous genes. (B) OrthoSNAP will identify five subgroups of single-copy orthologous genes (light blue) by identifying maximally inclusive subgroups—subtrees where each taxon is represented by a single sequence—and maximally inclusive subgroups after species-specific inparalog trimming (species-specific inparalogs are shown in orange). In contrast, DISCO and maximally inclusive strategies will identify three SC-OGs, in part, because they do not account for species-specific inparalogs. PhyloTreePruner, which only prunes species-specific inparalogs, will not identify any subgroups of single-copy orthologous genes due to the presence of more ancient duplication events.

**Supplementary table 1. Species and accessions for proteomes used in each dataset.** This table details the species used for the budding yeasts, filamentous fungi, and mammalian datasets. All proteomes from budding yeasts were downloaded from Shen et al. (2018) *Cell*. Proteomes from filamentous fungi and mammals were downloaded from NCBI and their accessions and assembly names are provided.

**Supplementary table 2. Number of orthogroups examined.** A table of the number of orthogroups, the number of SC-OGs, the number of gene families with orthologs and paralogs (MC-OGs), and the number of SNAP-OGs examined in the present study.

**Supplementary table 3. Ortholog occupancy for each dataset.** A table summarizing the average and standard deviation of taxon completeness in SC-OGs and SNAP-OGs.

**Supplementary table 4. Nine properties of phylogenetic information content.** Phylogenetic information content of SC-OGs and SNAP-OGs were examined using the nine properties described here. The abbreviation, description, additional notes, and function in PhyKIT used to calculate each property are listed here.

**Supplementary table 5. Multi-factor analysis of variance results reveals no substantial differences between SC-OGs and SNAP-OGs.** Degree of freedom, sum of squares, mean square, F-value and p-value for multi-factorial analysis of variance are shown here. Multi-factorial analysis of variance was conducting accounting for potential interaction effects as well as using an additive model, which does not account for interaction effects.

**Supplementary table 6. Tree certainty and tree certainty-all results.** Examining tree certainty and tree certainty-all revealed similar levels of incongruence among gene trees inferred using SC-OGs and SNAP-OGs.

**Supplementary table 7. Dataset for examining deep evolutionary relationships among eutherian mammals.** The NCBI accession, assembly name, name in files, and ingroup/outgroup designations are detailed here for each proteome used.

**Supplementary table 8. Number of orthogroups examined among eutherian mammals.** A table of the number of orthogroups, the number of SC-OGs, the number of gene families with orthologs and paralogs (MC-OGs), and the number of SNAP-OGs examined among eutherian mammals.

**Supplementary table 9. Gene support frequency results among ancient eutherian mammalian relationships.** Gene support frequency results reveal similar levels of support between the three hypotheses concerning deep evolutionary divergences among mammals. Multi-test corrected p-values are also shown here.

**Supplementary table 10. Comparison between different algorithms that identify subgroups of orthologous genes or conduct species-specific inparalog trimming.** Notably, OrthoSNAP provides the most user flexibility and handles the most use cases.
